# Accurate profiling of forensic autosomal STRs using the Oxford Nanopore Technologies MinION device

**DOI:** 10.1101/2021.07.01.450747

**Authors:** Courtney L. Hall, Rupesh K. Kesharwani, Nicole R. Phillips, John V. Planz, Fritz J. Sedlazeck, Roxanne R. Zascavage

## Abstract

The high variability characteristic of short tandem repeat (STR) markers is harnessed for human identification in forensic genetic analyses. Despite the power and reliability of current typing techniques, sequence-level information both within and around STRs are masked in the length-based profiles generated. Forensic STR typing using next generation sequencing (NGS) has therefore gained attention as an alternative to traditional capillary electrophoresis (CE) approaches. In this proof-of-principle study, we evaluate the forensic applicability of the newest and smallest NGS platform available – the Oxford Nanopore Technologies (ONT) MinION device. Although nanopore sequencing on the handheld MinION offers numerous advantages, including on-site sample processing, the relatively high error rate and lack of forensic-specific analysis software has prevented accurate profiling across STR panels in previous studies. Here we present STRspy, a streamlined method capable of producing length- and sequence-based STR allele designations from noisy, long-read data. To demonstrate the capabilities of STRspy, seven reference samples (female: n = 2; male: n = 5) were amplified at 15 and 30 PCR cycles using the Promega PowerSeq 46GY System and sequenced on the ONT MinION device in triplicate. Basecalled reads were processed with STRspy using a custom database containing alleles reported in the STRSeq BioProject NIST 1036 dataset. Resultant STR allele designations and flanking region single nucleotide polymorphism (SNP) calls were compared to the manufacturer-validated genotypes for each sample. STRspy generated robust and reliable genotypes across all autosomal STR loci amplified with 30 PCR cycles, achieving 100% concordance based on both length and sequence. Furthermore, we were able to identify flanking region SNPs with >90% accuracy. These results demonstrate that nanopore sequencing platforms are capable of revealing additional variation in and around STR loci depending on read coverage. As the first long-read platform-specific method to successfully profile the entire panel of autosomal STRs amplified by a commercially available multiplex, STRspy significantly increases the feasibility of nanopore sequencing in forensic applications.

## Introduction

Autosomal short tandem repeats (STRs) are the preferred genetic marker system for analyzing DNA evidence in forensic investigations. The high repeat length variability observed at STRs across the human genome facilitates individualization of evidentiary items and identification of the respective sources [1–3]. Current approaches to STR typing involve multiplex PCR amplification of forensically relevant loci followed by length-based separation and detection of fluorescently labeled amplicons using capillary electrophoresis (CE) [4–6]. The resultant STR profiles are therefore capable of resolving alleles of different repeat lengths but do not contain information regarding the underlying sequence composition of each allele. Although often sufficient for routine forensic casework, the discriminatory power achieved via CE may still be inadequate for more challenging samples even when additional autosomal STRs and other genetic markers are included alongside the 20 loci in the expanded core CODIS panel [7–10].

Advances in DNA sequencing technologies have enabled forensic researchers to ascertain nucleotide-level variation both in and around STRs with increasing ease [11–14]. The potential to harness all of the information contained within STR loci has led to a significant amount of interest in the use of next-generation sequencing (NGS) for human identification. NGS data has been used to differentiate between alleles of the same length but distinct sequence composition or motif organization (isoalleles), revealing up to two times as many alleles compared to CE [11]. Detection of hidden variation not only within, but also around STRs, along with the enhanced multiplex capabilities of NGS platforms over CE, greatly expands upon the resolution and discriminatory power of current STR panels [7,11,13,15].

Despite continued efforts focused on the development of streamlined, forensic-specific workflows [16,17] and data analysis software [18,19], widespread adoption of sequence-based STR typing has been hindered by the relatively high startup fees, involved workflows, and steep learning curves associated with NGS platforms. Moreover, the sequence-by-synthesis technique employed by Illumina platforms involves enzymatic reactions, fluorescently labeled nucleotides, and costly imaging equipment, in addition to multiple rounds of PCR for library enrichment and nucleotide determination [20,21]. The recent development and commercialization of nanopore sequencing devices by Oxford Nanopore Technologies (ONT) has brought the potential to bypass some of the major obstacles facing the application of NGS for forensic genetic analyses [20,22]. Nanopore sequencing relies on the translocation of molecules through nanopore proteins to determine the composition of nucleotides in native strands of DNA. Briefly, application of an electric voltage across a nanopore-containing membrane produces a constant ionic current through each of the pores within a given flow cell. Disruptions in the baseline current occur as individual strands of DNA are passed through the pore. These current disruptions, which are unique to the motif of three to five bases present in the pore, are recorded by the ONT device (e.g., MinION) and subsequently decoded to determine the sequence of nucleotides [23,24]. Nanopore sequencing platforms are therefore capable of directly sequencing reads of any length [25], eliminating some of the key biases associated with other NGS platforms [20,22].

Nanopore sequencing on the portable MinION device offers numerous advantages that could be particularly beneficial for processing fragile evidentiary material on-site at crime scenes. This unique sequencing technique is capable of simultaneously interrogating STRs on autosomal and sex chromosomes as well as other markers of forensic interest. Moreover, nanopore sequencing platforms are scalable to the output needs and financial restrictions of a given laboratory [20,26]. The cost of the handheld MinION is a small fraction of the initial investment required for implementation of other NGS platforms [21]. More recently, ONT has released the VolTRAX II device, which can facilitate implementation of a streamlined forensic workflow with minimal human intervention [20]. While the currently high cost of disposable flow cells and reagents would prohibit use in routine casework, this will likely decrease with increased commercial competition and further improvements in smaller-scale sequencers (e.g., Flongle) in the near future. Collectively, these features could position the ONT MinION device as an efficient and cost-effective alternative to mainstream NGS platforms for achieving the most comprehensive representation of genetic variability within a given sample. However, the relatively high error rate, particularly in homopolymers and low-complexity repeats [25,27,28], and lack of STR analysis software are significant obstacles to implementation of nanopore sequencing in forensics.

Few studies to date have assessed the use of the ONT MinION device for sequencing forensic STRs [29–32]. Despite successful profiling of the SNPs interrogated, only 14 of the 27 autosomal STRs were correctly typed across all samples using the most recently developed pipeline [29]. These and other researchers have attributed the inability to obtain complete and accurate STR profiles to the high error rate of ONT platforms [29–32]. The results obtained were used to identify locus- and allele-specific features (e.g., repeat number and motif complexity, presence of homopolymers) that prevent successful genotyping using nanopore sequencing data and provide guidelines for developing panels of ONT-compatible STR loci. Thus, a method that would enable us to expand upon the resolution and comprehensiveness of traditional approaches using established PCR multiplexes is still lacking and would be extremely beneficial.

In this study, we show that we can successfully resolve 22 autosomal STRs amplified with the Promega PowerSeq 46GY System and sequenced on the ONT MinION device (Fig. 1a). We developed and assessed a custom bioinformatics pipeline capable of producing accurate allele designations, including full STR and flanking region sequences, while accounting for long-read errors and PCR-induced stutter (Fig. 1b). The results obtained herein demonstrate that nanopore sequencing reads analyzed with our method (STRspy) can be used to achieve high-accuracy forensic STR profiles. Further, by capitalizing on sequence-level data, we can simultaneously detect variants in STRs and flanking regions separated by haplotype, allowing us to assess stutter as a consequence of PCR amplification across 15 and 30 cycles. Altogether, this is the first comprehensive approach to identify high-resolution genomic variants in ONT sequencing data for forensic STR typing purposes.

**Fig. 1.**
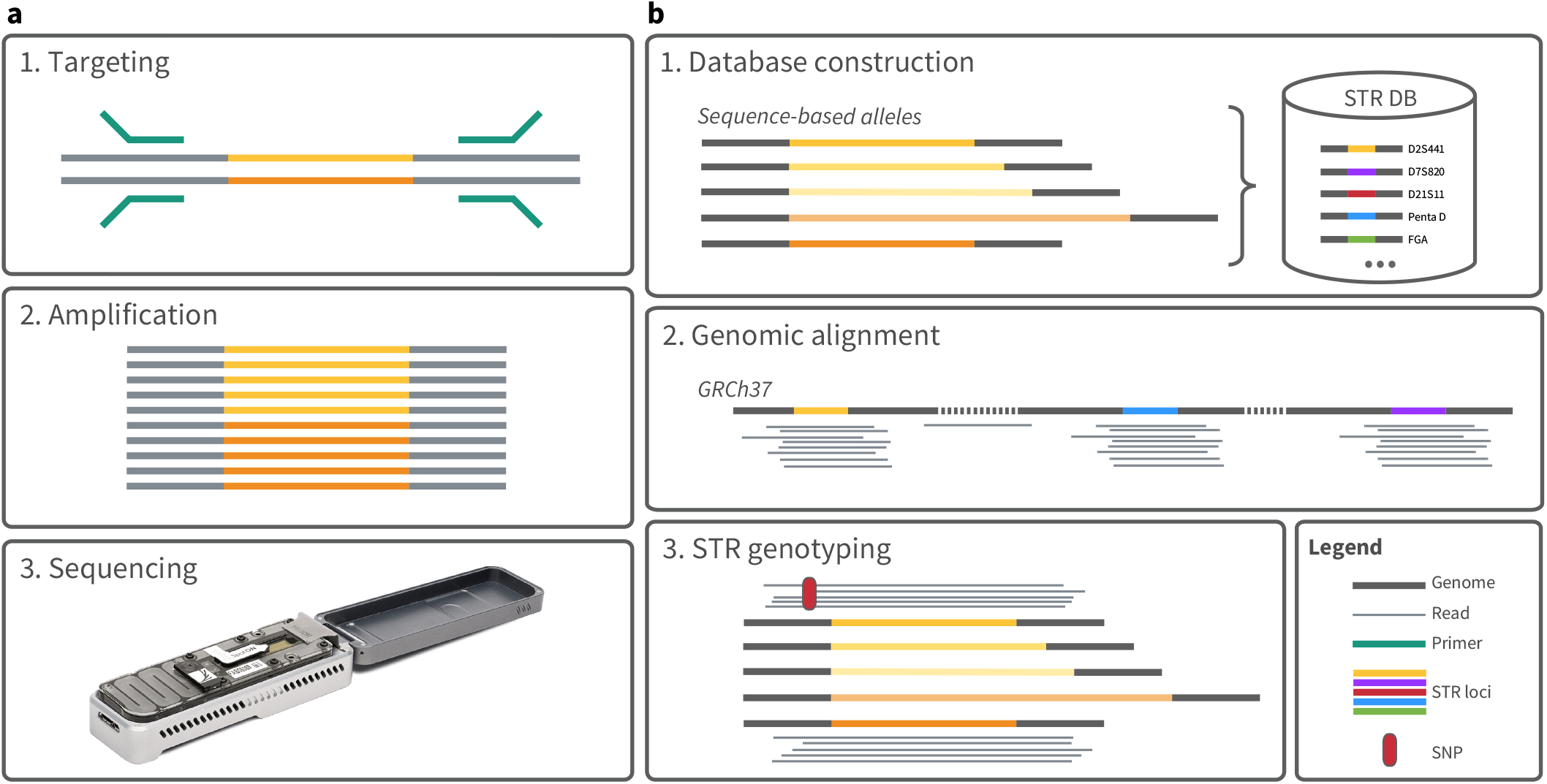
STR sequencing and profiling with STRspy. a) Lab workflow. STR loci are targeted and amplified via multiplex PCR. Amplicon libraries are then prepared and sequenced on the ONT MinION device to generate nucleotide-level data. b) Data analysis pipeline. STRspy relies on a user-generated STR database (DB) containing sequence-based alleles for each locus of interest. Reads are first aligned to the human reference genome. Reads overlapping STR loci are then extracted and mapped to the custom STR DB. STRspy uses the normalized read counts to rank the STR alleles and predict the genotype at each locus. Sequencing data produced and analyzed as described can resolve alleles of the same length but different underlying sequence (dark yellow and orange) and identify SNPs in the flanking region (red). See figure legend for more details.

## Materials & Methods

### Samples

The results presented in this paper are based on sequencing data from six NIST traceable standards and one Promega control (female n = 2; male n = 5). Extracted DNA along with validated length- and sequence-based genotype information for these reference samples were obtained directly from the respective manufacturers. Promega single-source male DNA 2800M for human STR analysis was normalized to 0.1ng/μL based on the manufacturer-specified quantification value. Components A, B, and C of NIST Standard Reference Material (SRM) versions 2391c and 2391d were quantified on the Qubit 2.0 Fluorometer using the Qubit dsDNA BR Assay Kit (Thermo Fisher Scientific) and diluted to a concentration of 0.1ng/μL. The same methods were used to verify the final concentration of all samples prior to downstream applications.

### PowerSeq amplification

Six full PCR reactions per sample were prepared with the Promega PowerSeq 46GY System (PS4600) according to the manufacturer’s technical manual with 0.5ng input DNA. Amplification was performed in triplicate on an Eppendorf Mastercycler pro S using the recommended thermal cycling conditions at either 15 or 30 cycles. Resultant amplicons were then subject to an Agencourt AMPure XP bead (Breckman Coulter) cleanup (2.5:1 ratio based on sample volume) to remove remaining primers and PCR reaction components.

### Nanopore library preparation & sequencing

Purified PCR products were multiplexed and prepared for nanopore sequencing using the ONT Ligation Sequencing Kit (SQK-LSK109) with Native Barcoding Expansion 1-12 (EXP-NBD104). Library preparation was performed with the following modifications to the standard Native Barcoding Amplicons protocol (NBA_9093_v109_revC_12Nov2019). Amplicon DNA input used for library preparation fell below the recommended 1μg for all samples. Quantification steps were conducted on the Agilent TapeStation 4200 with D1000 ScreenTape for samples amplified at 30 cycles. To minimize pore clogging and maximize yield of the short amplicon libraries, no more than 75ng was loaded onto each individual flowcell (based on previous optimizations studies; data not shown). Following ligation of ONT sequencing adapters, the 30-cycle pooled barcodes were quantified and diluted to 75ng in elution buffer (EB, ONT) if necessary. Quantification steps were completely bypassed for 15-cycle amplicon libraries, and the entire volume of barcoded samples were combined. To further reduce potential sample loss due to bead purification, pooled barcodes exceeding the volume required for subsequent steps (>65μL) were concentrated in an Eppendorf 5301 Vacufuge System. Prepared libraries were loaded in a drop-wise fashion into the SpotON port of primed vR9.4D flow cells (FLO-MIN106D, ONT). Flow cells were placed in the MinION device and sequenced until exhaustion (up to 72 hours) using the ONT MinKNOW software. Details regarding sample multiplexing are provided in Supplementary Table S1.

### Bioinformatics pipeline & algorithm description

#### Implementation

STRspy is designed to predict forensic STR genotypes from long-read sequencing data. STRspy requires a minimum of one thread and is executed at the command line. We implemented and tested this framework in a Unix/Linux environment. STRspy is under MIT license (open source) and can be downloaded from GitHub (https://github.com/unique379r/strspy). The GitHub page also includes associated documentation and a small test set to verify successful installation.

STRspy relies on a user-generated reference database to produce allele designations consistent with the established forensic naming system [33]. The same STR database can be used to analyze any samples of interest, and thus users are only required to build it once. We constructed the database for this study using STR sequencing data for 1036 samples published under the STRSeq BioProject (NIST 1036) [12]. Our STR database includes all reported sequence-based alleles for the 22 PowerSeq autosomal loci along with 500bp flanks. Each entry is labeled with the locus name, bracketed repeat motif, and length-based allele designation used in standard STR profiling (Supplementary Fig. S1). The custom STR database produced in this study is available at https://github.com/unique379r/strspy.

STRspy accepts basecalled reads in the form of either fastq or bam files to accommodate both ONT and PacBio data. Users are also required to provide bed and fasta files for the STR database (see below). STRspy executes the following three steps in a per sample manner (Supplementary Fig. S2):

1. Basecalled reads are first aligned to the human reference genome (hg19/GRCh37) with minimap2 (v2.18-r1015) [34]. STRspy includes predefined parameters to adapt minimap2 to either ONT or PacBio read data. Subsequently, the mapped reads are automatically converted and sorted into a bam file using samtools (v1.12) [35].
2. The genome-wide bam file is processed with bedtools intersect (v2.30.0) [36] to extract reads that overlap STR loci of interest based on the locations specified in the user-provided bed file. The extracted locus-specific reads are then mapped to the predefined collection of alleles contained within the custom STR database using minimap2 (v2.18-r1015) [34]. As in the previous step, STRspy generates sorted bam files containing the mapped reads.
3. STRspy computes the number of reads (with mapping quality greater than 1) mapped to each sequence-based STR allele in the sorted bam files with samtools (v1.12) [35]. This part of the pipeline can be implemented in a multi-threaded manner to increase the speed of analysis. STRspy calculates locus-specific normalized read counts by dividing the number of reads per allele across the highest number of reads mapping to a single allele at each STR. Both the raw and normalized read counts are stored for subsequent filtering and assessment of the results. STRspy uses the normalized read counts to rank the STR alleles at each locus and reports either a single allele (homozygous) or the top two alleles (heterozygous) based on the user-defined normalization threshold. By default, this threshold is set to 0.4.

#### SNP detection

STRspy uses xAtlas [37] to detect SNPs within the flanking regions of each autosomal locus contained within the STR database and region bed file. SNP calls produced by xAtlas are output in vcf file format which is compatible with various available bioinformatic tools for downstream data analysis. We filtered resultant vcf files to keep SNP calls with "PASS" flags and p-values of 0.8 or higher. To prevent the accumulation of incorrect SNP calls due to differences in sequencing depth [38], samples amplified at 30 PCR cycles were uniformly subsampled to 1% of total mapped reads with samtools view -s 0.01 (v1.12) [35]. The randomly subsampled datasets were then used for SNP calling and benchmarking of the 30-cycle dataset.

### Data analysis

Raw signal data collected on the MinION device were basecalled and separated by barcode with the standalone version of Guppy (v3.4.2). Merged fastq files from the seven samples amplified at 15 and 30 PCR cycles in triplicate were processed using the STRspy command line interface to obtain normalized read counts, length- and sequence-based allele designations, and SNP calls in the flanking regions. The utility scripts available on the STRspy GitHub repository (https://github.com/unique379r/strspy) were implemented to assess the overall performance of STRspy, evaluate concordance between predicted and known genotypes, identify stutter artifacts, and visualize results as heatmaps and line plots.

Manufacturer-validated genotypes obtained via CE and NGS served as the ground truth for assessing STRspy performance based on error rate calculations. Correct allele predictions produced by STRspy were classified as true positives, incorrect as false positives, and dropout as false negatives. Precision and recall were calculated as the correct STR allele designations (true positive) out of total alleles reported by STRspy (true positive + false positive) or the ground truth dataset (true positive + false negative), respectively. F1 score, which provides a measure of overall test accuracy, was determined by taking the harmonic mean of precision and recall. These metrics were calculated with normalization cutoffs ranging from 0.1 to 0.9 to identify the optimal threshold at both cycle numbers (Fig. 2a, Supplementary Fig. S3). Allele designations obtained at this cutoff (0.4) were used as the STRspy predictions for overall performance assessments (Fig. 2b).

**Figure 2.**
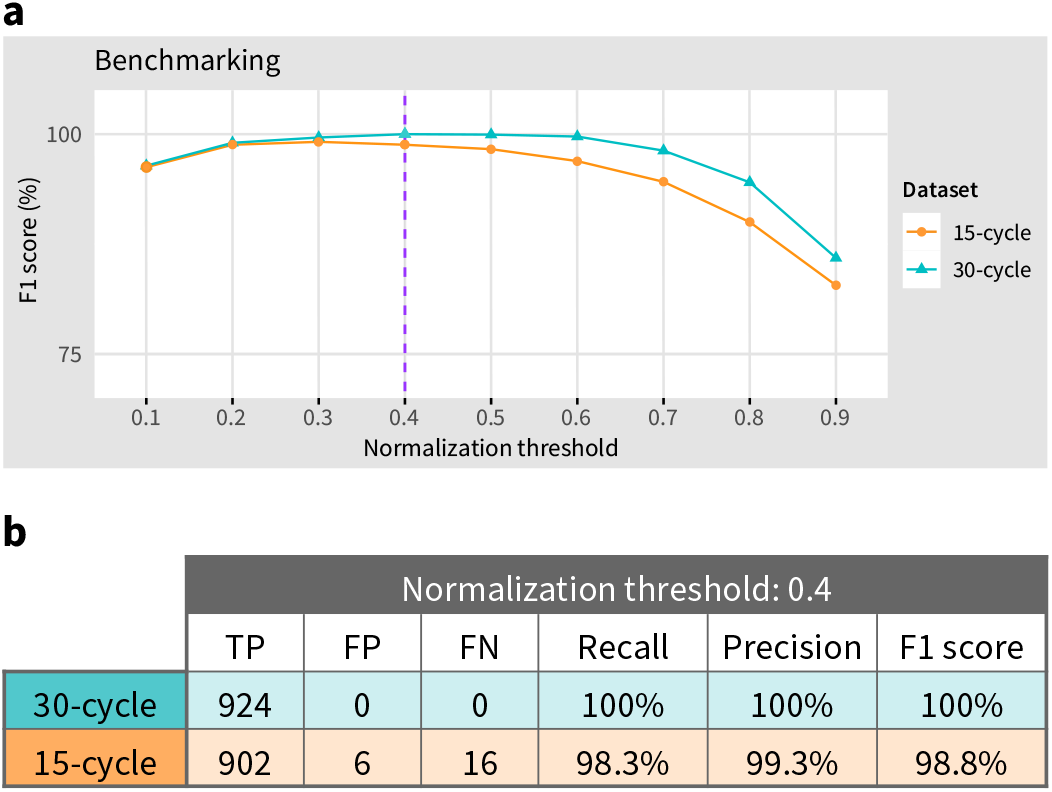
STR benchmarking. a) Plot of F1 score across different normalization thresholds. b) Table showing the number of true positive (TP), false positive (FP), and false negative (FN) predictions produced by STRspy as well as associated benchmarking metrics at the normalization threshold used in this study (0.4).

## Results

### Assessing forensic STR loci on the ONT MinION device

As a relatively new sequencing platform, the ONT MinION device has undergone limited testing for forensic DNA analyses. To assess the capabilities of this device in the context of human identification, we processed seven reference samples with validated length- and sequence-based STR profiles (see methods). In short, 22 autosomal STRs were amplified at 15 and 30 PCR cycles using the Promega PowerSeq 46GY System with 0.5ng of input DNA and sequenced on the MinION. Each sample was processed in triplicate at both cycle numbers, resulting in ONT sequencing data for 42 individual amplicon libraries. To analyze the nucleotide-level data produced, we developed STRspy, a novel method for the detection and characterization of forensic STR loci using ONT and PacBio reads (see methods). STRspy not only identifies different STR alleles present at a given locus but does so in a phased manner. Moreover, STRspy is often able to detect SNPs present in the flanking region, thus leveraging all information contained within the amplicons.

As expected, the number of reads produced for each sample varied based on PCR cycle number (Supplementary Table S2). The percent of total reads that mapped to STR loci for samples in the 30-cycle dataset ranged from 87.76% to 92.87% with an average of 90.76%. More variability was observed across the 15-cycle dataset, in which on-target efficiencies fell between 50.96% and 71.67% and averaged 65.09%. Nonetheless, the raw read counts mapped to STR loci were comparable across 15-cycle samples. Similarly, depth of coverage per locus was impacted by PCR cycle number, resulting in a mean of 246,002.27 and 321.56 reads in the 30- and 15-cycle datasets, respectively. We also observed PCR amplification bias that resulted in reduced – and sometimes insufficient – coverage over several loci, particularly D22S1045 (Supplementary Table S3). The effect of amplification bias on genotype determination was overcome with increased PCR cycles. Overall, these data suggest that PCR amplification followed by nanopore sequencing results in high on-target rates and enables in-depth analysis of allelic content.

We assessed STRspy performance at two distinct amplification cycle numbers in triplicate samples, providing novel insight into the level of coverage required for accurate genotype determination and reproducibility. Length- and sequence-based allele designations as well as flanking region SNP calls were generated for samples in each dataset using ONT sequencing reads and STRspy (see methods). STRspy employs a user-defined threshold to predict if a locus is heterozygous (reporting the top two alleles) or homozygous (reporting the top allele) based on the normalized coverage supporting each STR allele. To determine the optimal cutoff value, we assessed STRspy performance at different normalization thresholds (Fig. 2a, Supplementary Fig. S3). We also measured the runtime of STRspy using a single thread for each sample. The average runtime across samples in the 30-cycle dataset was 571 minutes (9.51 hours) due to the high depth of coverage (mean: 246,002.27). We observed a significant reduction in STRspy runtime for the lower coverage 15-cycle dataset (mean: 321.56), which averaged 3.54 minutes per sample. STRspy is implemented to support multithreading and thus runtimes can be improved using multiple CPU cores to increase analysis speed.

### Length- & sequence-based genotype determinations

We assessed the true positive (i.e., correct STR allele), false positive (i.e., incorrect STR allele or additional STR allele at known homozygous loci), and false negative (i.e., missing STR allele at heterozygous loci) rates for each autosomal STR compared to the manufacturer-validated genotypes. Using these metrics, we were able to determine if STRspy fully recovered known STR genotypes and correctly assigned allele designations for each individual sample.

STRspy was able to consistently predict the correct allele designations based on both length and sequence for all 22 autosomal loci amplified at 30 PCR cycles (Fig. 3b). The utility of ONT sequencing data analyzed with STRspy is demonstrated by the 30-cycle triplicates for NIST A from SRM 2391c (NISTAc). STRspy successfully identified repeats characterized by simple motifs such as the D2S441 tetranucleotide [TCTA]10 allele. Further, our method was able to resolve the length-based homozygous 10 alleles observed at this locus to produce heterozygous calls consisting of the simple [TCTA]10 and compound [TCTA]8 TCTG [TCTA]1 repeats (Table 1). Similar results were achieved for NIST B from SRM 2391d (NISTBd), which possesses isoalleles at DS2441 (11, 11). These data also enabled differentiation of isoalleles between samples, further increasing profile resolution.

**Table 1.** STRspy resolves isoalleles. Normalized read counts, raw read counts (parentheses), and STRspy predictions (bold) for isoalleles at D2S441 loci with repeat lengths of 10 or 11. Reported values are for triplicate 1 in the 30- and 15-cycle datasets.

| Repeat length | 30-cycle |  |  | 15-cycle |  |  |
| --- | --- | --- | --- | --- | --- | --- |
|  | [TCTA]10 | [TCTA]8 TCTG TCTA | Prediction | [TCTA]10 | [TCTA]8 TCTG TCTA | Prediction |
| **10** |  |  |  |  |  |  |
| 2800M | **0.681 (40203)** | 0.015 (861) | TP | **0.825 (70)** | 0.012 (1) | TP |
| NISTAc | **0.862 (72250)** | **1.0 (83848)** | TP | **0.984 (60)** | **1.0 (61)** | TP |
| NISTBc | **0.737 (88101)** | 0.057 (6768) | TP | **0.853 (64)** | 0.027 (2) | TP |
| NISTCc | 0.035 (7280) | **1.0 (206237)** | TP | 0.032 (3) | **1.0 (95)** | TP |
| **11** |  |  |  |  |  |  |
|  | [TCTA]11 | [TCTA]9 TCTG TCTA | Prediction | [TCTA]11 | [TCTA]9 TCTG TCTA | Prediction |
| NISTAd | **1.0 (83390)** | 0.022 (1851) | TP | **1.0 (123)** | 0.016 (2) | TP |
| NISTBd | **1.0 (57604)** | **0.912 (52511)** | TP | **1.0 (80)** | **0.975 (78)** | TP |
| NISTCd | **0.993 (47865)** | 0.036 (1710) | TP | **0.906 (106)** | 0.009 (1) | TP |

**Fig. 3.**
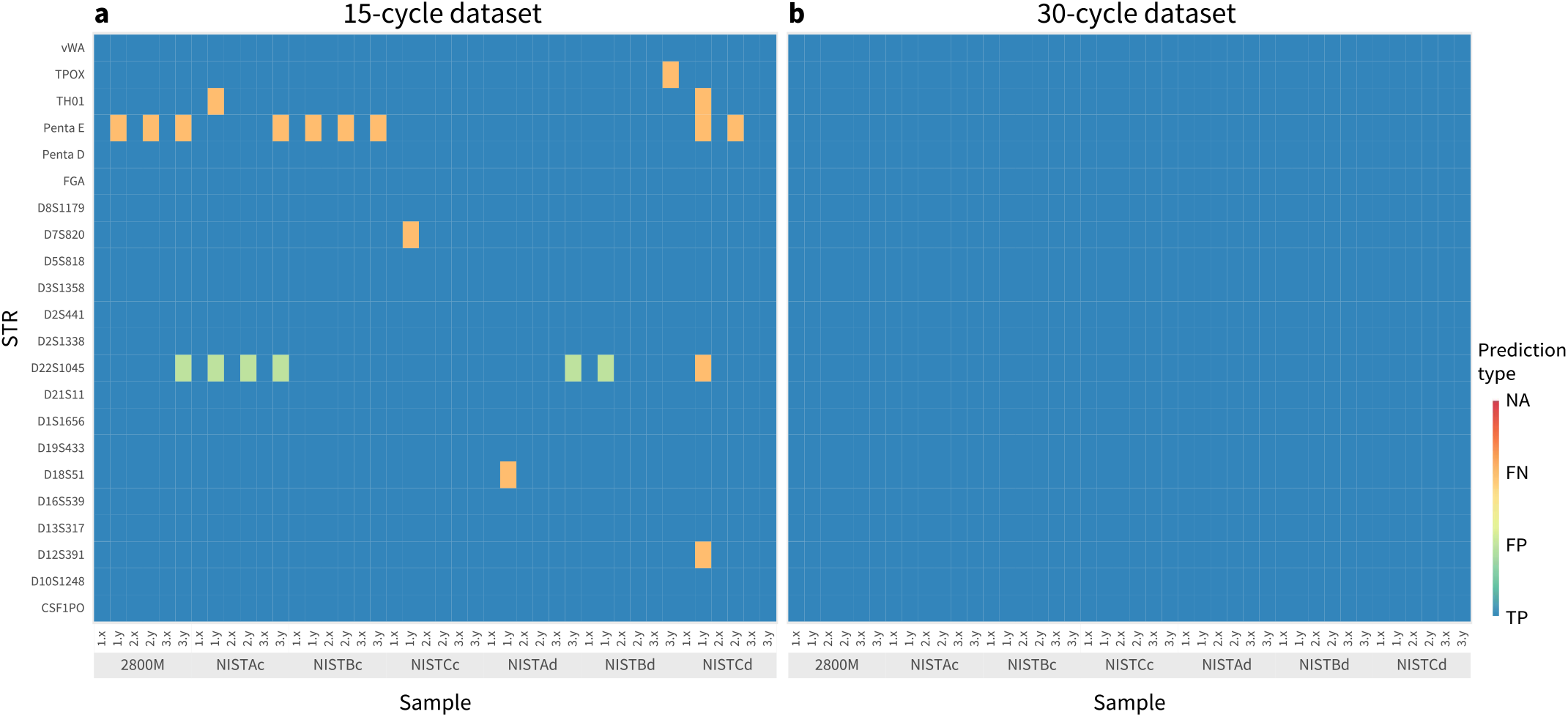
STR allele designations. Heatmap comparison of STRspy predictions to manufacturer-verified length- and sequence-based genotypes across the a) 15-cycle dataset and b) 30-cycle dataset. True positive predictions are depicted in blue, false positives in green, and false negatives in orange. Reference samples (grey boxes) are labeled by triplicate (1, 2, 3) and haplotype (x, y).

Despite variation in raw and normalized read counts, STRspy was able to resolve sequence-based heterozygous alleles of the same length using ONT reads across the 22 loci. Complete concordance was achieved for all samples amplified at 30 PCR cycles, resulting in 100% recall, precision, and F1 score (Fig. 2b). These observations demonstrate the ability of our method to (1) differentiate alleles of the same length but different sequence and (2) accurately genotype simple, compound, and complex repeat motifs using ONT sequencing data.

Next, we evaluated the ability of STRspy to profile the same seven samples at 15 PCR cycles (Fig. 3a). STR loci amplified with a lower number of PCR cycles had less coverage compared to those in the 30-cycle dataset (Supplementary Table S2). Nevertheless, STRspy distinguished between the length-based homozygous 10 and 11 alleles at D2S441, predicting the correct heterozygote genotypes across all three 15-cycle triplicates for samples in this dataset (Table 1). Other repeat motifs are composed of homopolymers and intervening sequences that are not counted toward the length-based genotypes produced via CE. Even with this low level of coverage, STRspy predicted the correct allele designations at the homopolymer-containing Penta D and complex FGA loci. Allele designations concordant with CE were also obtained for D19S433 and D21S11 despite the presence of intervening sequences that complicate length-based profiling from sequencing data.

Precision, recall, and F1 score for the 15-cycle datasets were 99.34%, 98.26%, and 98.80%, respectively (Fig. 2b). The 22 incorrect genotypes (out of 924) produced by STRspy fall into two distinct categories of errors: false positives (i.e., two alleles predicted at a known homozygous locus, or the incorrect allele predicted) and false negatives (i.e., one allele predicted at a known heterozygous loci). All six of the false positive allele designations were observed at D22S1045 due to the relatively low coverage over this locus (Supplementary Table S3) and the presence of stutter artifacts (see below). The other 16 errors were false negatives, an overwhelming majority (9) of which were at Penta E (Fig. 4a). False negatives at this particular locus across all samples were characterized by allele dropout of the longer repeating unit. For instance, in one NISTAc triplicate, STRspy correctly predicted [AAAGA]5 but not [AAAGA]10 at Penta E (5, 10). Although a greater number of raw reads supported the 10 allele for the false negative genotype (15-cycle.3: 99 reads) compared to a true positive genotype (15-cycle.1: 38 reads), the normalized read count for the 15-cycle.3 NISTAc triplicate fell below the 0.4 threshold.

**Fig. 4.**
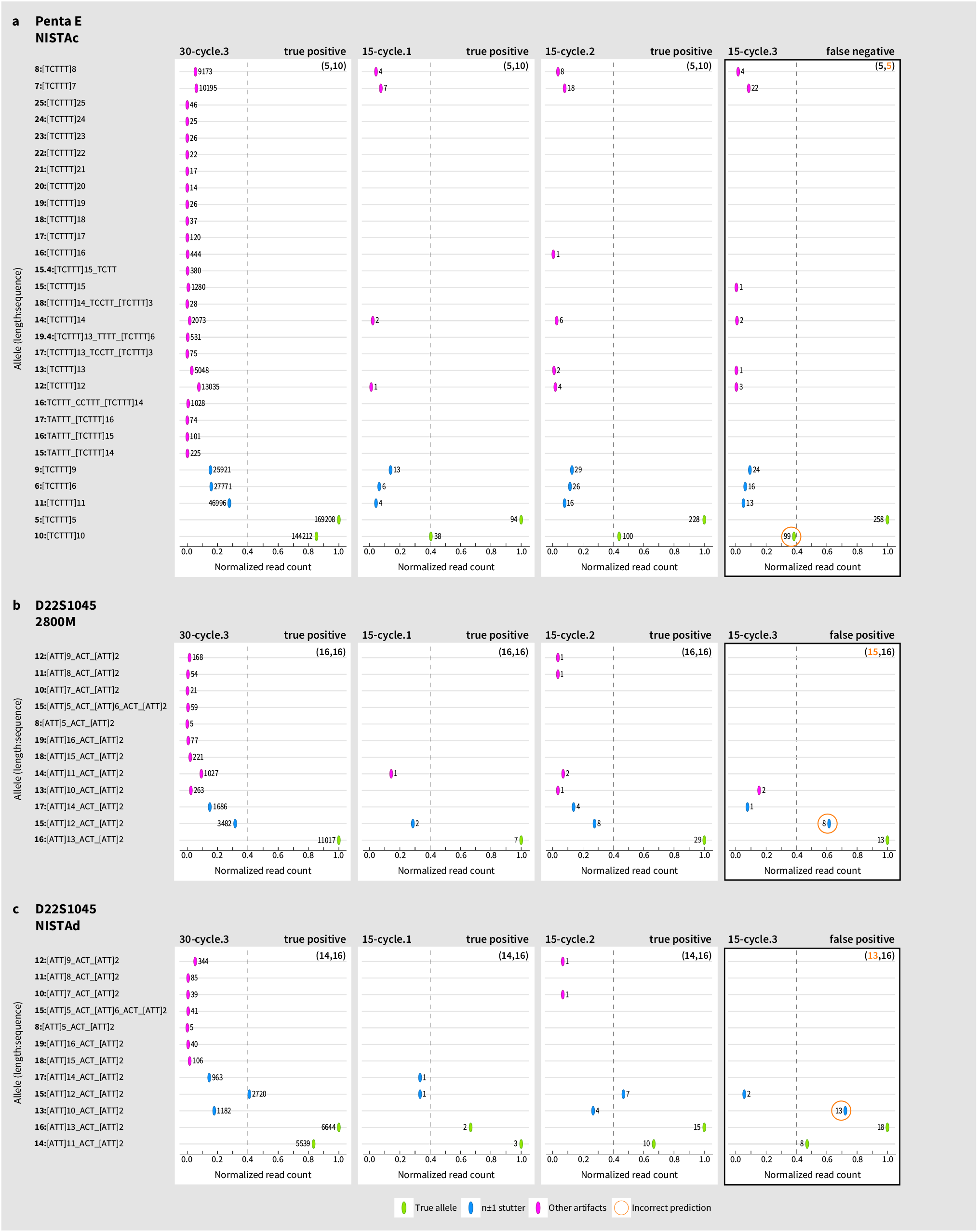
Genotyping errors. a) False negative genotype due to allele drop out at Penta E for NISTAc. b) False positive genotype due to stutter artifacts at D22S1045 for the b) 2800M homozygote and c) NISTAd heterozygote. Raw read counts are included next to each point. The incorrectly typed triplicate in each set is denoted by a black box and the incorrect allele prediction is circled in orange. One 30-cycle (left) and all three 15-cycle triplicates are shown for comparison purposes. See figure legend for more details.

In contrast to Penta E, STRspy correctly predicted the longer [AATG]6 ATG [AATG]3 but not the shorter [AAGT]8 for TH01 in one of the NISTAc triplicates. Further examination of the individual NISTAc datasets in which these loci were problematic revealed a minor allele normalized count of 0.38 and 0.31 for Penta E and TH01, respectively (Supplementary File S1). Consequently, decreasing the normalization cutoff value to 0.3 increased the 15-cycle F1 score from 98.80% to 99.13% by preventing minor allele dropout (Supplementary Fig. S3). These observations ultimately suggest that the prevalence of false negatives is due to amplification bias and lack of locus coverage rather than inherent limitations of STRspy itself.

### Flanking region variation

Single nucleotide polymorphisms (SNPs) as well as insertions and deletions (indels) have been observed in sequences around forensic STRs. These variants further increase the discriminatory power of current STR panels but cannot be detected in the length-based profiles generated via CE. We therefore examined the ability of STRspy to detect known flanking region SNPs in the NIST SRMc/d samples. Detailed benchmarking results are provided in Table 2.

**Table 2.** SNP benchmarking. Comparison of filtered calls generated by xAtlas to known flanking region SNPs in the 30- and 15-cycle NIST triplicates. DB position is the SNP position in our STR database with respect to the associated STR allele and 500bp flanks. 30-cycle data were subsampled as described in the main text.

| Sample | Locus | STR | SNP | DB position | dbSNP ID | 30-cycle |  |  | 15-cycle |  |  |
| --- | --- | --- | --- | --- | --- | --- | --- | --- | --- | --- | --- |
|  |  |  |  |  |  | Recall | Precision | F1 score | Recall | Precision | F1 score |
| NISTCc | D13S317 | 11 | T | 545 | rs9546005 | 66.67% | 100% | 80% | 100% | 50% | 66.67% |
| NISTAd | D16S539 | 13 | C | 406 | rs1728369 | 100% | 42.86% | 60% | 100% | 50% | 66.67% |
|  | D1S1656 | 15.3 | T | 569 | rs4847015 | 100% | 100% | 100% | 100% | 100% | 100% |
|  | D1S1656 | 18.3 | T | 581 | rs4847015 | 100% | 100% | 100% | 100% | 100% | 100% |
|  | D5S818 | 11 | G | 557 | rs25768 | 100% | 42.86% | 60% | 100% | 100% | 100% |
|  | D7S820 | 8 | A | 479 | rs7789995 | 100% | 100% | 100% | 100% | 100% | 100% |
| NISTBd | D13S317 | 11 | T | 545 | rs9546005 | 33.33% | 16.67% | 22.22% | 100% | 100% | 100% |
|  | D16S539 | 9 | C | 552 | rs11642858 | 100% | 100% | 100% | 100% | 100% | 100% |
|  | D16S539 | 11 | C | 406 | rs1728369 | 66.67% | 50% | 57.14% | 66.67% | 40% | 50% |
|  | D1S1656 | 15.3 | T | 569 | rs4847015 | 100% | 100% | 100% | 100% | 100% | 100% |
|  | D2S1338 | 17 | A | 466 | rs6736691 | 100% | 100% | 100% | 100% | 75% | 85.71% |
|  | D5S818 | 12 | A | 497 | rs73801920 | 100% | 37.50% | 54.55% | 100% | 50% | 66.67% |
|  |  |  | G | 561 | rs25768 |  |  |  |  |  |  |
|  | D5S818 | 12 | G | 561 | rs25768 | 100% | 27.27% | 42.86% | 100% | 50% | 66.67% |
|  | D7S820 | 10 | A | 479 | rs7789995 | 100% | 100% | 100% | 100% | 100% | 100% |
| NISTCd | D13S317 | 14 | T | 557 | rs9546005 | 100% | 42.86% | 60% | 100% | 100% | 100% |
|  | D16S539 | 9 | C | 552 | rs11642858 | 66.67% | 100% | 80% | 100% | 100% | 100% |
|  | D5S818 | 13 | G | 565 | rs25768 | 100% | 75% | 85.71% | 100% | 100% | 100% |
|  | D5S818 | 15 | G | 573 | rs25768 | 100% | 60% | 75% | 100% | 100% | 100% |
|  | D7S820 | 9 | A | 479 | rs7789995 | 100% | 60% | 75% | 100% | 50% | 66.67% |
|  |  |  | A | 545 | rs16887642 |  |  |  |  |  |  |
|  | D7S820 | 10 | A | 479 | rs7789995 | 100% | 100% | 100% | 100% | 100% | 100% |
|  | TPOX | 8 | A | 405 | rs145426142 | 100% | 100% | 100% | 100% | 100% | 100% |
| Overall |  |  |  |  |  | 92.06% | 74.05% | 82.08% | 98.41% | 84.05% | 90.66% |

We first assessed the SNP calls at samples amplified with 30 PCR cycles. Poor performance was observed for the non-subsampled 30-cycle dataset, indicating that excessive coverage (mean: 246,002.27) hinders SNP calling due to the accumulation of sequencing errors. For this reason, reads were subsampled and reanalyzed as in previous publications [38]. The recall and precision achieved by the subsampled 30-cycle dataset were 92.06% and 74.05%, respectively. The reduced coverage (mean: 321.56) obtained with fewer amplification cycles in the 15-cycle dataset eliminated the need for subsampling. We recovered known SNPs across all but one of the samples amplified with 15 PCR cycles (rs1728369 in NISTB.1 from SRM 239c). The overall recall and precision for the 15-cycle dataset were 98.41% and 84.05%, respectively. These results show that lower coverage actually improves our ability to identify SNPs in terms of both precision and recall.

Unlike SNPs, indels in the flanking region impact the length-based allele designations used in forensics. Flanking region indels were therefore incorporated into the STR database itself and can be identified by inspecting the bracketed repeat motif reported by STRspy. For instance, a subset of alleles observed at D13S317 are characterized by a rare 4bp deletion in the flanking region [12]. Consequently, the length-based 11 allele can correspond to [TATC]11 or [TATC]12. The latter repeat motif, in which the 4bp deletion occurs within the 3’ flank, is identical to that of a 12 length-based allele but is identified as an 11 via CE. Despite these complexities, STRspy was able to distinguish between the [TATC]12 with the 4bp flanking region deletion (2800M), [TATC]12 (NISTBc, NISTAd, NISTCd), and [TATC]11 (NISTCc, NISTBd) to produce the correct sequence- and length-based genotypes a across all samples at this locus.

### Impact of amplification cycle number on stutter artifacts

Polymerase slippage during amplification of low-complexity repeats can lead to stutter artifacts in resultant datasets [39]. In contrast to the sequence-by-synthesis technique harnessed by Illumina platforms, nanopore sequencing relies on the direct detection of nucleotides in each strand of DNA [21]. This unique capability provides novel insight into PCR-induced bias. To assess the impact of amplification cycle number on stutter artifact formation we examined the prevalence of reads one repeat unit smaller and larger than the true allele (n±1) with the STRspy utility scripts (Supplementary File S1). Previous STR genotyping attempts have been complicated by the presence of stutter artifacts at D18S51 ([AGAA]n) in ONT sequencing data [29]. Consistent with the notion that stutter percentage increases with the number of PCR cycles, we observed higher normalized read counts for n±1 stutter at D18S51 when NISTAc (12, 15) was amplified with 30 cycles (0.36, 0.38, 0.41) compared to 15 cycles (0.26, 0.28, 0.27). Nevertheless, STRspy was able to identify the correct alleles at both cycle numbers even when the normalized read count for stutter exceeded 0.4 (as in Fig. 4c: 30-cycle.1 and 15-cycle.2).

We also investigated how stutter artifacts contribute to the false positive results in the 15-cycle dataset. As mentioned, all six false positive allele designations were observed at D22S1045. The raw count data (Supplementary Table S3) revealed that a relatively low number of reads mapped to this locus across samples in both datasets, which is indicative of amplification bias. Consequently, STRspy called an additional allele at D22S1045 for one of the 2800M samples amplified with 15 PCR cycles (Fig. 4b). The majority of reads mapped to the true homozygous allele (16) at this locus. However, the presence of stutter in the minus direction (15) exceeded the normalization threshold of 0.4 and was therefore called by STRspy. Similar observations were made at one of the NISTAd triplicates (Fig. 4c). In contrast to the allele drop-in for 2800M (which is homozygous at D22S1045), STRspy produced the incorrect designations for the shorter alleles in the NISTAd (14, 16) heterozygote. In the 30-cycle.3 and 15-cycle.2, the 15 allele (representing the overlap of minus stutter for the 16 allele and plus stutter for the 14 allele) exceeds the normalization threshold but falls below the true alleles in rank and thus is not reported by STRspy. Interestingly, the incorrect allele prediction for 15-cycle.3 was minus stutter (13) associated with the minor allele (14) rather than the overlap (15). Collectively, these observations highlight the stochastic nature of PCR-induced artifacts as well as the impact of amplification bias on genotyping errors.

## Discussion

In this paper, we report the first STR analysis for accurate predictions of allele designations along with identification of flanking region SNPs specific to long-read sequencing platforms. Using the Promega PowerSeq Kit and the ONT MinION device we produced robust sequencing data across all targeted loci that enabled us to investigate the impact of PCR cycle number. Although a higher number of reads mapped to PCR artifacts in the 30-cycle dataset, the normalized read counts for the true alleles exceeded stutter, resulting in the correct predictions at all loci. The 15-cycle dataset was skewed due the low level of coverage, and thus the 30-cycle dataset produced more reliable genotypes. Additionally, we showcased the accurate identification of STR alleles and flanking regions SNPs. These results suggest that this portable, scalable, and rapid sequencing approach could prove extremely valuable in future applications. A maximum of 4 samples were pooled and sequenced on a single MinION flow cell. Given the high level of coverage achieved at 30 PCR cycles, it may be possible to increase the number of samples sequenced on a single MinION flow cell (without exceeding 75ng total) to reduce overall cost. STRspy leverages ONT sequencing data to profile STRs with unprecedented accuracy regardless of repeat motif, complexity, or length. All relevant studies to date have reported incorrect genotypes at vWA, FGA, and D21S11 due to repeat pattern complexity [29–32]. We demonstrated that the novel method developed and tested in the current paper was able to produce the correct length- and sequence-based allele designations for vWA, FGA, and D21S11 across all samples even at the low-level coverage obtained from 15 PCR cycles.

While the length and continuous sequence information obtained in this study enabled accurate STR identification and phasing using STRspy, current long-read ONT data still suffers from certain biases (e.g., homopolymer error rates) that may impact the performance of STRspy. We predict that recent and future developments from ONT (e.g., in the base calling algorithm) will improve the quality of sequencing data, further increasing the accuracy and performance of STRspy-based analyses. Despite current biases in ONT sequencing data, STRspy predicted the correct genotypes for the homopolymer-containing Penta D and Penta E using 30-cycle reads produced on the standard R9 nanopore proteins. Although Penta D was also correctly typed across all samples in the 15-cycle dataset, dropout of the minor Penta E allele was observed in nine samples. Reducing the normalization cutoff value from 0.4 to 0.3 resulted in the correct genotype, suggesting that the establishment of locus-specific normalization thresholds in future studies may be beneficial when analyzing low-coverage samples. Given the improvements in STR data quality previously reported [30], the use of the R10 nanopore proteins may further mitigate this issue by increasing the number of usable reads produced from lower cycle numbers.

In addition to producing robust and reliable STR profiles, STRspy possesses numerous features that support implementation in forensic genetics without the need for extensive bioinformatic training in long-read data processing and analysis. Our easy-to-install method can be used on computational infrastructures ranging from personal laptops to high-performance clusters, closely mirroring the scalability of nanopore sequencing platforms. Furthermore, STRspy executes all steps required to go from basecalled reads to STR profiles based on user-defined parameters and input files. The minimal computational requirements and streamlined nature of STRspy not only increases the overall accessibility of ONT sequencing in forensic genetics, but also supports field applications. Although beyond the scope of the current study, samples processed with the ONT Field Sequencing Kit should be analyzed with STRspy to establish protocols using the limited laboratory and computational equipment that would be available at crime scenes.

The ability of STRspy to achieve correct genotype predictions depends on the alleles being present in the STR database provided by the user. Because the custom reference database we generated contains the most common STR alleles observed among the four major U.S. populations [12], it can be used to profile many unknown samples. This list however is not exhaustive, and the database constructed for this study only includes autosomal STRs amplified by the Promega PowerSeq 46GY System. Future efforts will be geared towards adding the 23 PowerSeq Y-STRs to our database and assessing profiles generated using STRspy. Users can also expand upon or create their own database containing STRs of interest if sequence-based allele information is available and formatted in the same manner across all loci.

The data analyzed herein was produced in a conventional laboratory setting using high-quality DNA extracts. Each sample was amplified and sequenced in triplicate, providing novel insight into the reproducibility of STR profiling on the MinION device. Although we sequenced more amplicon libraries than all relevant publications [29–32] our study included a small number of unique samples. Additional experiments involving more reference and probative samples will be conducted in future studies. These data will allow us to evaluate STR profiling from biological material of similar quality to that collected from suspects and crime scenes, respectively. Further assessment of amplification at various cycle numbers and input DNA concentrations will also provide important information about the sensitivity of nanopore sequencing devices and expand upon our understanding of PCR-induced artifacts. These experiments will ultimately form the foundation for establishing ONT-specific protocols and interpretation guidelines for STR profiling.

A key limitation of sequence-based STR typing in routine forensic casework is cost. Despite the relatively low startup fee of the ONT MinION device, the price per sample exceeds that of mainstay short-read sequencing platforms [26]. The rapid evolution of nanopore sequencing since the 2014 release of the MinION device has led to a significant decrease in error rate and increase in throughput that have, in turn, reduced overall cost [27,40]. Continued technological improvements and developments (e.g., Flongle adapter and flow cells) in coming years will likely reduce the cost and increase the accessibility of ONT sequencing. Despite the cost of sequencing, the ONT MinION device provides unique advantages including higher resolution over current typing techniques and faster turnaround time with potential on-site analyses. These features would be particularly beneficial in forensic investigations.

## Conclusion

This study assessed the ability to profile forensic STRs using nanopore sequencing data produced on the ONT MinION device. With our novel forensic-specific analysis method, STRspy, we were able to achieve robust and reliable STR profiles for all autosomal loci amplified at 30 cycles with the Promega PowerSeq 46GY System. The results presented herein demonstrate that nanopore sequencing platforms are capable of producing length-based allele designations consistent with standard forensic nomenclature while revealing an additional level of variation in and around STR loci. The novel pipeline we developed overcomes the issues reported in previous publications to profile the entire panel rather than a subset of STRs amplified by a commercially available kit. We anticipate that continued improvements in nanopore sequencing technologies, along with further development of STRspy, will increase the feasibility of forensic STR profiling on ONT devices in not only a traditional laboratory setting, but also on-site at crime scenes.

## Acknowledgments

We would like to thank the Budowle lab and Promega for providing the validated sequence-based allele designations for the 2800M reference sample.

