## Supplementary Fig. S1-S3 for "Accurate profiling of forensic autosomal STRs using the Oxford Nanopore Technologies MinION device"

length-based designation

bracketed motif

locus

**sequence-based allele #1**

```
>D2S441_[TCTA]10_10
GGTTTCCAAGATCCCATTTGGGTTTGATTAATTTGCCAGCGTTGCTCACAGAAATCAAGG
AAACACTTATTTAACGTCTGCTGGTTTGTTATAAATAATGTTACAAAGGATAACAATGAA
GAGATGGTCAGGCGAGGTATGGGGGAAGGGGCGTGGAGCTTCCATGTCCTCCCTGGGCGC
CACCCTCCAGGAACCTCCACGTGTTCAGCTATACAGAAGCTTCCTGAACCCAGTCCTCTT
GGGGTTTGAGGGAAGCTTCATGACATCAGCATTCCTTCCTCCAGGGTATTAATGGGACCC
TCTCTGAAGAGATTCTTAAGACCCACGGCCAGAAAGTTGGGTAAAGACTAGAGTCCTGCC
TTGGGGCAGGTGAAAGGAGTGCAAGAGAAGGTAAGAGAGATTCTGTTCCTGAGCCCTAAT
GCACCCAACATTCTAACAAAAGGCTGTAACAAGGGCTACAGGAATCATGAGCCAGGAACT
GTGGCTCATCTATGAAAACTTCTATCTATCTATCTATCTATCTATCTATCTATCTATCTA
TATCATAACACCACAGCCACTTAGCTCCAATTTAAAAGATTAATCATAAACATTTGGGAA
GGAGAGTGAAGATTTTTGTGATGTTAAATAAGAATGATTATACTAAAAACCAAAATAATA
TGTTATTTATGGCTGGGTGTGGTGGCTTAAGCCTGTAATCCCAGAACTTTGGGAGGCCAA
GGCTTGTGGATCACTTGAGCCCAGAAGTTCAAGACCAGCCTGGGCAACATAGGGAGACCC
TGTCTCTACAAAAAATTTTAAAATTAGCTGGACATGATGGCACGCACCCGTAGTCTCAGC
TACTCAGGAGGCTCACGCCACTGCATTCCAGTCTGGGTAACGCACACCTTATCTCTAAAA
AATATGTATGTGTGTGTACATATATATATACATATATATACATACATATATATACATATA
TACATACATATATACATATATACACATATATACATATAATACACACATATTTACATATAT
ACACACATATATACATATAT
```

STR

500bp flanks

**sequence-based allele #2**

```
>D2S441_[TCTA]8_TCTG_TCTA_10
GGTTTCCAAGATCCCATTTGGGTTTGATTAATTTGCCAGCGTTGCTCACAGAAATCAAGG
AAACACTTATTTAACGTCTGCTGGTTTGTTATAAATAATGTTACAAAGGATAACAATGAA
GAGATGGTCAGGCGAGGTATGGGGGAAGGGGCGTGGAGCTTCCATGTCCTCCCTGGGCGC
CACCCTCCAGGAACCTCCACGTGTTCAGCTATACAGAAGCTTCCTGAACCCAGTCCTCTT
GGGGTTTGAGGGAAGCTTCATGACATCAGCATTCCTTCCTCCAGGGTATTAATGGGACCC
TCTCTGAAGAGATTCTTAAGACCCACGGCCAGAAAGTTGGGTAAAGACTAGAGTCCTGCC
TTGGGGCAGGTGAAAGGAGTGCAAGAGAAGGTAAGAGAGATTCTGTTCCTGAGCCCTAAT
GCACCCAACATTCTAACAAAAGGCTGTAACAAGGGCTACAGGAATCATGAGCCAGGAACT
GTGGCTCATCTATGAAAACTTCTATCTATCTATCTATCTATCTATCTATCTATCTGTCTA
TATCATAACACCACAGCCACTTAGCTCCAATTTAAAAGATTAATCATAAACATTTGGGAA
GGAGAGTGAAGATTTTTGTGATGTTAAATAAGAATGATTATACTAAAAACCAAAATAATA
TGTTATTTATGGCTGGGTGTGGTGGCTTAAGCCTGTAATCCCAGAACTTTGGGAGGCCAA
GGCTTGTGGATCACTTGAGCCCAGAAGTTCAAGACCAGCCTGGGCAACATAGGGAGACCC
TGTCTCTACAAAAAATTTTAAAATTAGCTGGACATGATGGCACGCACCCGTAGTCTCAGC
TACTCAGGAGGCTCACGCCACTGCATTCCAGTCTGGGTAACGCACACCTTATCTCTAAAA
AATATGTATGTGTGTGTACATATATATATACATATATATACATACATATATATACATATA
TACATACATATATACATATATACACATATATACATATAATACACACATATTTACATATAT
ACACACATATATACATATAT
```

**Supplementary Fig. S1.** STR database structure. The STR database required by STRspy consists of the text string of nucleotides corresponding to each sequence-based allele (yellow) along with 500bp flanks (orange). Each entry is labeled with the locus name (pink), bracketed repeat motif (blue), and length-based allele designation used in forensics (green). The isoalleles with a repeat length of 10 at D2S441 are depicted.

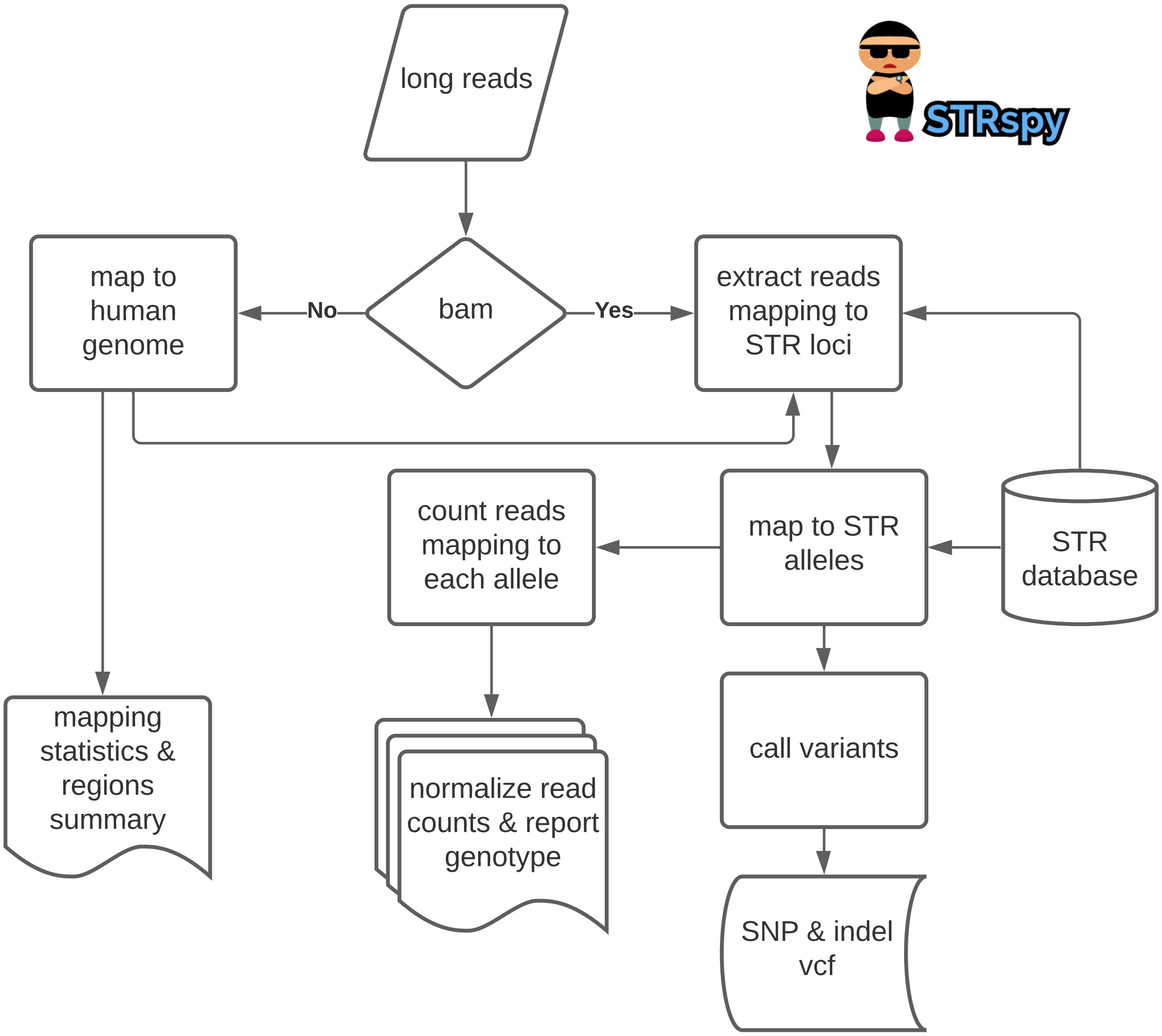


**Supplementary Fig. S2.** Steps implemented in STRspy. See main text for additional details.

**a**

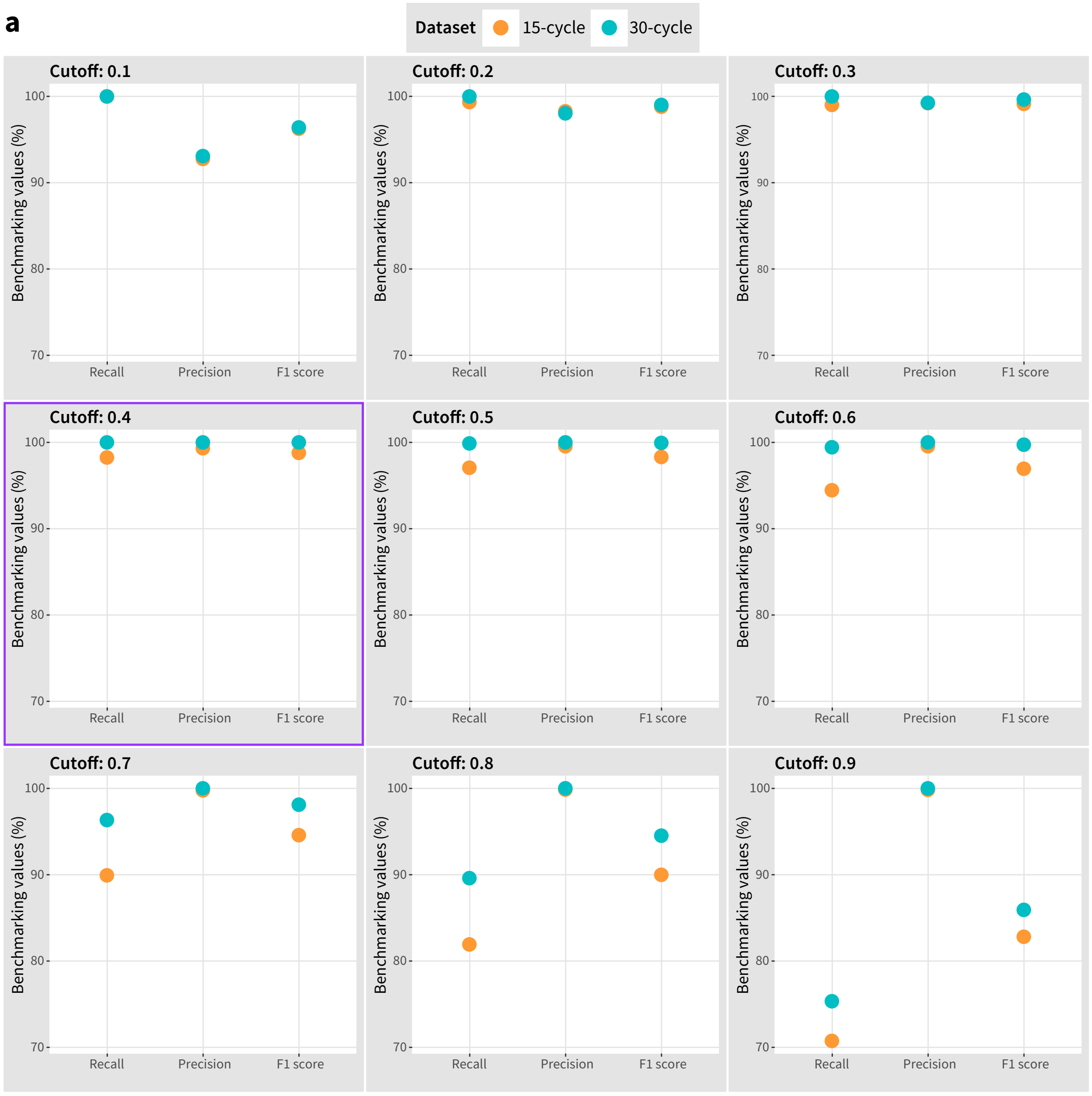


**b**

| | 30-cycle | | | 15-cycle | | |
|---|---|---|---|---|---|---|
| **Cutoff** | **TP** | **FP** | **FN** | **TP** | **FP** | **FN** |
| **0.1** | 860 | 64 | 0 | 857 | 67 | 0 |
| **0.2** | 906 | 18 | 0 | 902 | 16 | 6 |
| **0.3** | 917 | 7 | 0 | 908 | 7 | 9 |
| **0.4** | 924 | 0 | 0 | 902 | 6 | 16 |
| **0.5** | 923 | 0 | 1 | 893 | 4 | 27 |
| **0.6** | 919 | 0 | 5 | 869 | 4 | 51 |
| **0.7** | 890 | 0 | 34 | 829 | 2 | 93 |
| **0.8** | 828 | 0 | 96 | 756 | 1 | 167 |
| **0.9** | 696 | 0 | 228 | 653 | 1 | 270 |

**Supplementary Fig. S3.** a) Benchmarking plots across a range of normalization thresholds for the 30- and 15-cycle datasets. b) Table showing the number of true positive (TP), false positive (FP), and false negative (FN) predictions produced by STRspy at each cutoff tested. The optimal cutoff (0.4) and default normalization threshold for STRspy is denoted by purple boxes.
